# Neolithic super-grandfather Y haplotypes, their related surnames, and autism spectrum disorder

**DOI:** 10.1101/077222

**Authors:** Pei He, Na Chen, Zhengmao Hu, Zuobin Zhu, Kun Xia, Shi Huang

**Affiliations:** Laboratory of Medical Genetics, School of Life Sciences, Central South University 110 Xiangya Road, Changsha, Hunan, 410078, China

**Author notes:** Present address: Department of Genetics, Xuzhou Medical University, Xuzhou, Jiangsu 221004, China (Zuobin Zhu).

**Keywords:** Neolithic super-grandfather Y haplotypes, Yan and Huang, assortative mating, social economic status (SES), autism spectrum disorder (ASD)

## Abstract

Recent studies found three Neolithic super-grandfather Y haplotypes among Han Chinese, consistent with the legend of Yan and Huang Emperors. Individuals of royal and noble ancestry or high social economic status (SES) are known to practice assortative mating and consanguineous marriages, which can produce offspring of both higher and lower than average fitness. However, the roles of the super-grandfather Y haplotypes and their descendant lines in history, fitness, and the male biased autism spectrum disorder (ASD) remain unknown. Here we show a link between the super-grandfathers and the legend of *Yan-Huang* Emperors and between their descendant haplotypes and ASD. We found that subjects carrying the O3a1c and O3a2c1a super-grandfather haplotypes were enriched with *Yan* and *Huang* related surnames, respectively, in two independent datasets of 1564 and 772 male Han subjects. We identified high and low SES descendant haplotypes of the super-grandfathers using the Han dataset of the 1000 genomes project based on two criteria: more descendant branches and fewer mutations before star-like expansions. By genotyping 505 fathers of ASD affected male children from the Autism Clinical and Genetic Resources in China with surnames either closely related to Huang (Ying group) or less related (Ji group), we found the high SES haplotypes within the O3a2c1a clade at ∼2 fold lower (odds ratio 2.05, 95% CI 1.28-3.26, P=0.0026) while the low SES haplotypes at ∼2 fold higher frequency (odds ratio 1.92, 95% CI 1.01-3.64, P = 0.046) in the fathers relative to 505 normal subjects. The fraction of low SES haplotypes was greater than that of high SES in ASD fathers of the Ying group, in contrast to Ying controls or the Ji fathers and Ji controls. Consistently, analysis of 2366 ASD affected children showed higher male to female ratio for Ying versus Ji group (6.52 +/-1.11 v 4.59 +/-0.41, P = 0.028, one tailed). These results provide evidence for the Yan-Huang legend and suggest a role for Y in ASD.

## Introduction

The Han Chinese population uses hereditary surnames that are thought to be first established ∼5000 years ago (*1-3*). Modern Han Chinese people are thought to be largely descended from *Yan Di* (Yan Emperor) and *Huang Di* (*Huang* Emperor) who lived ∼5000 years ago. There were “Eight Great *Xings* of High Antiquity” from ∼4000 years ago that are believed to be ancestors of most of today’s ∼23813 surnames of Chinese people (*4*). Although these Eight Great *Xings* are thought to originate in matriarchal societies, it is expected that certain males may be more dominant than others in such societies. *Yan* belonged to one of the Eight Great *Xings* (Jiang) and the other 7 Great *Xings* are all related to *Huang*. Of these, Ji is thought to be the original surname of *Huang* and has the most descendant surnames today. Ying is special because one of its related contemporary surnames, Huang, is also the same as the commonly used name for the Yellow Emperor or *Huang Di*.

Consistent with Neolithic individuals matching the legendary status of *Yan*-*Huang*, there were three Neolithic super-grandfathers (*5*). Their Y haplotypes originated ∼5.4 Kya (thousand years ago) for O3a2c1a-Page23 or M117 (O2a2b1a1, ISOGG 2017), ∼6.5 Kya for O3a2c1-F46 (O2a2b1a2a1), and ∼6.8 Kya for O3a1c-F11 (O2a1c1a1a,), and represent 16%, 11%, and 14% of present Han Chinese, respectively. Based on the estimated age and frequency, O3a2c1a-Page23 could be a good candidate for *Huang* and O3a1c-F11 for *Yan*. Therefore, we here tested whether contemporary Han males with surnames or *Xings* more closely related to *Yan* and *Huang* are enriched with O3a1c and O3a2c1a, respectively.

Individuals of royal and noble ancestry or high social economic status (SES) are known to practice assortative mating and consanguineous marriages (*6, 7*), which can produce offspring of both higher and lower than average fitness (*8*). One of the diseases associated with assortative mating is autism spectrum disorders (ASD) (*9-11*). Parents with ASD children often have mild forms of autistic-like characteristics (*12*). The male to female ratio is 4:1 in the global ASD population, but is 23:1 in ASD subjects without physical or brain abnormalities (*13*). Little is known about the male bias in ASD (*14*). We hypothesized that descendant haplotypes of a super-grandfather may show dimorphism, with some associated with high SES and better fitness while others the opposite. In conjunction with studying high SES haplotypes and surnames among normal Han Chinese subjects, we studied a large ASD cohort from the Autism Clinical and Genetic Resources in China (ACGC) (*15*).

## Results

We first made use of the Y haplotype data of surnames representing 1564 males as reported on the Website“One Surname a Week” maintained by researchers from Fudan University. To determine Y haplotype distribution among the Eight Great *Xings*, we divided contemporary surnames into 4 groups of Great *Xings*, Jiang, Ying, Ji, and Others according to popular surnames literatures (Supplementary Table S1). We obtained the average fraction of individuals per surname for each of the 3 super-grandfather haplotypes (Figure 1A). The Jiang-group has more O3a1c than each of the other groups (P < 0.05, Student’s t test, one tailed). The Ying-group has more O3a2c1a than each of the other groups (P < 0.02, Student’s t test, one tailed). To verify the above result, we collected peripheral blood samples from healthy subjects in the Hunan area in China and did PCR-sequencing on the 3 haplotypes. The results on 772 males again showed similar patterns of O3a1c enrichment in the Jiang-group, and O3a2c1a enrichment in the Ying-group (Figure 1B and Supplementary Table S2). The combined results from these two surveys were significant, as the probability of getting both haplotypes correctly matched to their respective candidate groups is 1/144 or 0.007 (the chance of randomly matching a haplotype to its surname group is 1/16 [4 groups and 2 surveys] and the chance of getting a second haplotype correctly matched is 1/9 [3 remaining groups and 2 surveys]).

**Figure 1.**
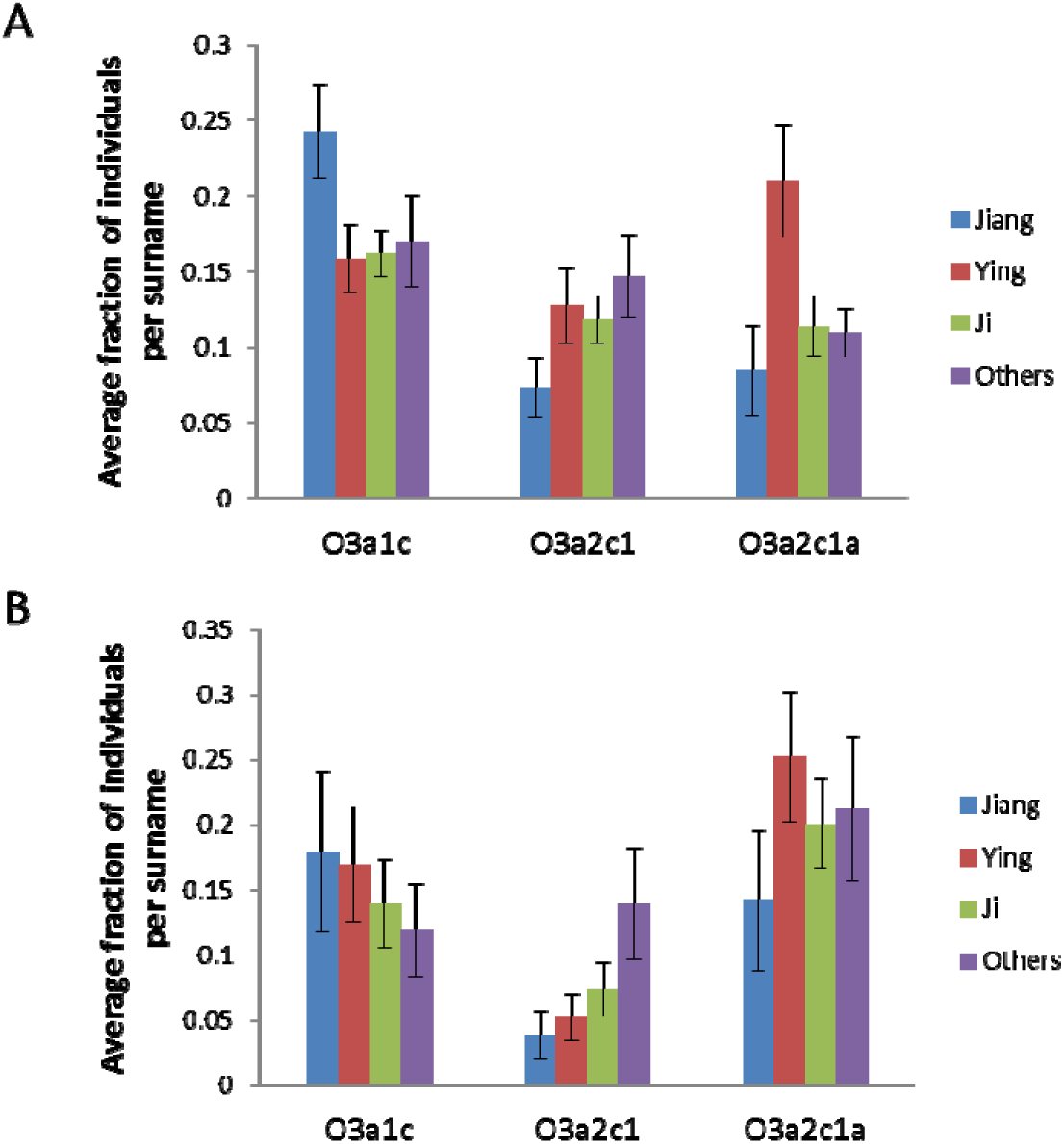
Distribution of the three super-grandfather Y haplotypes. The fraction of each haplotype in a surname was calculated and the average fractions per surname with standard error of the mean are shown in the plot. Shown are results from 1564 male subjects with surname and Y data from the website "One Surname a Week" (A) and from 772 male subjects collected in this study (B).

We made use a previous study on the 1000 genomes project (1kGP) dataset that showed star like expansion for the three super-grandfathers (Supplementary Figure S1) (*16*). We focused on Han Chinese in this dataset and assigned high SES Y haplotypes based on two criteria: more descendant branches and fewer mutations before star-like expansions since a super-grandfather’s early descendants were more likely to be high SES individuals. Within the O3a2c1a-Page23 clade, branch 225 as indexed by Poznik et. al. (2016) (Supplementary Figure S1) was identified as the super-grandfather haplotype (*16*). Under branch 225, 4 branches (168, 178, 185, and 188) appear to be high SES as each of these has only 1 SNP before further branching splits with branch 178 having the largest number of descendant lines. The remaining 3 branches (161, 219, and 222) were identified as low SES haplotypes among Han Chinese.

Within the O3a2c1-M1561 clade, the super-grandfather haplotype was identified as branch 150 defined by F46, which has 2 sub-branches with the sub-branch 149 as the more super-grandfather like. Of the 3 descendant lines from branch 149, the high SES branch was identified as 131, followed by 149 and 148. Within the O3a1c-002611 clade, the super-grandfather haplotype was identified as branch 69 defined by F11, which also has 2 sub-branches with the sub-branch 68 as the more super-grandfather like. Of the descendant lines from branch 68, the high SES branch was identified as 57, followed by 64 and 40.

To study the roles of Y and surnames in ASD, we focused on the O3a2c1a clade and the Huang related surname groups Ying and Ji. We genotyped 91 Ying and 414 (or 355 for some haplotypes) Ji normal subjects and 202 Ying and 303 Ji fathers of ASD affected male children (Supplementary Table S3). No significant differences in frequencies were observed between the normal and ASD subjects in the O3a2c1a-page23 haplotype (Figure 2). However, relative to normal Ying subjects, Ying fathers showed >3 fold lower fraction of the two high SES haplotypes 178 and 188 (P < 0.05). Relative to normal Ji subjects, Ji fathers showed 1.75 fold lower fraction of 178 (P< 0.05). As the low SES haplotypes each had too few samples to be informative, we combined Ying and Ji subjects and low or high SES haplotypes for further analysis. We found the high SES haplotypes (178, 188, 168 and 185 combined) at ∼2 fold lower (Odds ratio 2.05, 95% CI 1.28-3.26, P=0.0026) while the low SES haplotypes (222, 219 and 161) at ∼2 fold higher frequency (odds ratio 1.92, 95% CI 1.01-3.64, P = 0.046) in the fathers relative to normal Ying and Ji subjects (Figure 2B). The results indicated a relationship between ASD and Y haplotypes, especially for the Ying subjects relative to the Ji subjects, consistent with the above noted association between Ying and the O3a2c1a super-grandfather.

**Figure 2.**
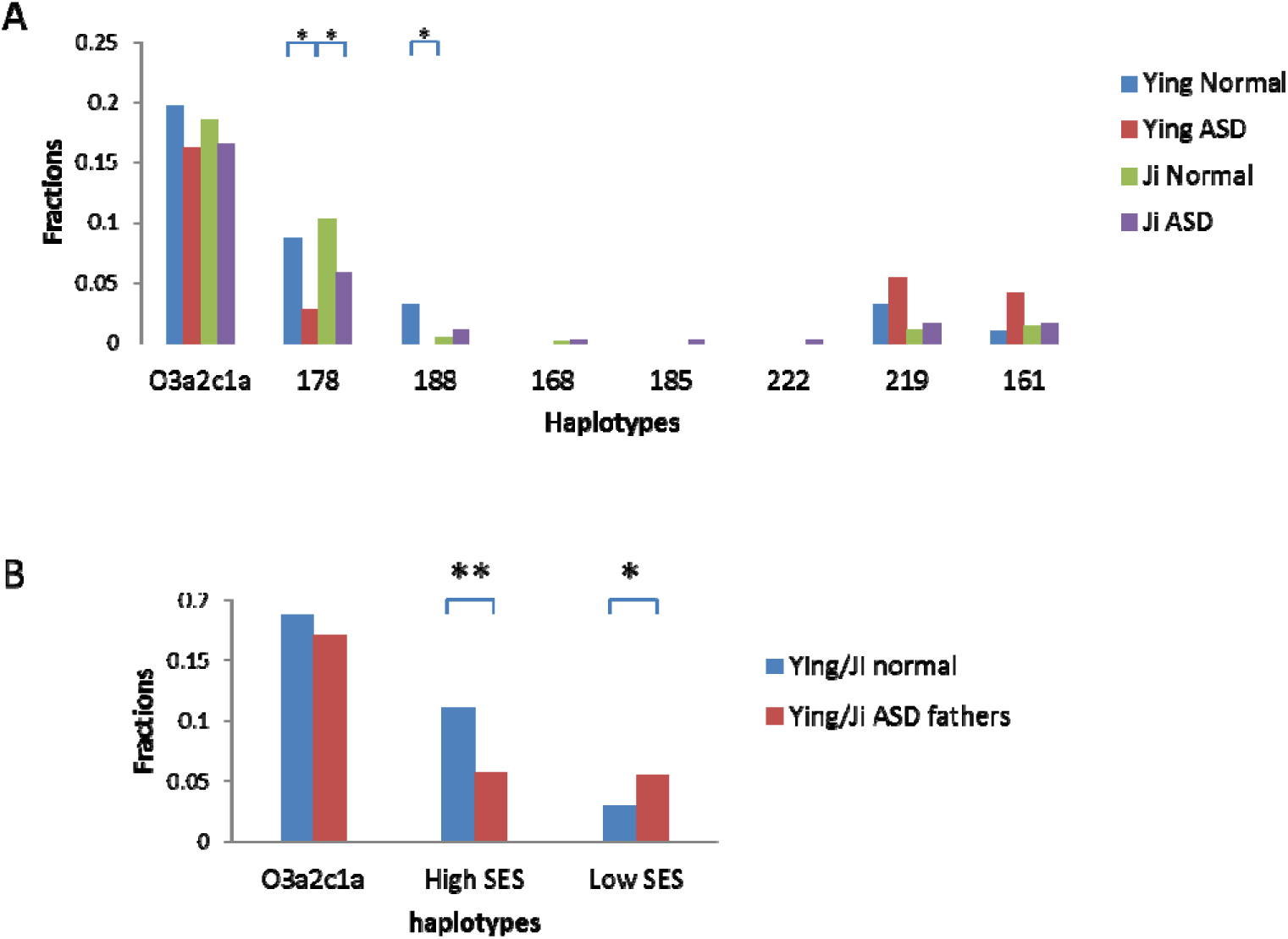
Frequency of O3a2c1a related haplotypes. **A.** Fractions of O3a2c1a and related downstream haplotypes in normal or fathers of ASD affected male children with Ying or Ji related surnames. **B.** Fractions of O3a2c1a and related downstream high SES (178, 188, 168 and 185) and low SES haplotypes (222, 219 and 161) in subjects of combined Ying and Ji groups. **, P < 0.01, *, P < 0.05, Chi square test, 2 tailed.

To confirm this pattern of ASD link with Y SES status, we next studied the other two super-grandfather haplotypes in the combined Ying and Ji subjects. For the O3a1c-002611 clade, ASD fathers showed 1.76 fold (P < 0.03, one tailed Chi square test) lower frequency in the high SES haplotype (branch 57) but no significant difference in the low SES haplotypes (branches 64 and 40) or the O3a1c-002611 haplotype relative to normal Ying and Ji subjects (Figure 3A). For the O3a2c1-M1561 clade, ASD fathers showed 2.22 fold (P <0.05, Chi square test, 2 tailed) and 1.52 fold (P <0.05, Chi square test, 2 tailed) higher frequency for the low SES haplotypes (branches 148 and 140) and the O3a2c1-M1561 haplotype, respectively (Figure 3B). These results provide additional data for a consistent link between ASD and SES status of Y haplotypes and ASD.

**Figure 3.**
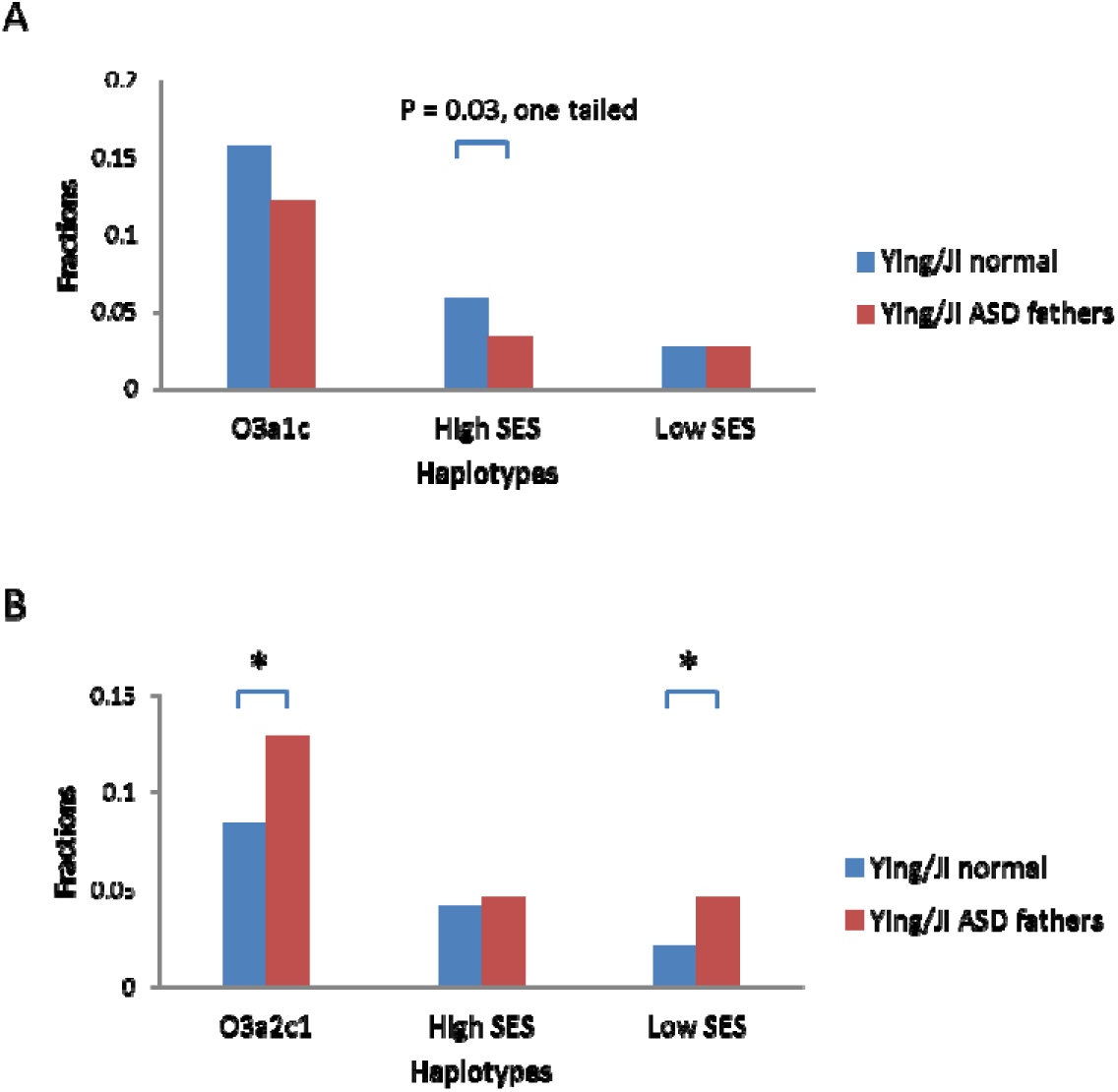
Frequency of O3a1c and O3a2c1 related haplotypes. **A.** Fractions of O3a1c and related downstream high SES (57) and low SES haplotypes (64 and 40) in subjects of combined Ying and Ji groups. **B.** Fractions of O3a2c1 and related downstream high SES (131) and low SES haplotypes (148 and 140) in subjects of combined Ying and Ji groups. *, P < 0.05, Chi square test, 2 tailed.

If certain low SES Y haplotypes are enriched in ASD, one would expect surnames linked with those haplotypes to be also enriched in ASD. The above analysis showed a more consistent enrichment of low SES Y haplotypes in ASD for the Ying relative to the Ji group (Figure 2A). We further confirmed this by finding that, while the majority of normal Ying subjects carried the top ranked SES haplotypes 178, 188, 168, 185, 131, and 57, most Ying ASD cases carried low SES haplotypes 161, 219, 148, 140, 64, and 40 (fractions of high and low SES were 0.25 and 0.12 for Ying normal and 0.11 and 0.17 for Ying ASD, respectively), which was in contrast to the Ji group (0.2 and 0.07 for Ji normal and 0.16 and 0.1 for Ji ASD, respectively). Therefore one expects ASD cases with Ying surnames to be more affected by Y and show more extreme male bias than Ji-related cases. We analyzed 2362 ASD affected children for their distribution among the top 40 surnames in China (with at least 2 female cases in our dataset here) and found 3 of 6 Ying-related surnames (Huang, Xu and Ma) ranked among the top 10 surnames in male to female ratio whereas only 1/13 Ji-related surnames did (P < 0.05, Chi square test, one tailed, Figure 4A). The average ratio of Ying group was higher than the other three groups and significantly higher than the Ji group (*P* < 0.05, Student’s t test, one tailed, Figure 4B, Supplementary Table S4).

**Figure 4.**
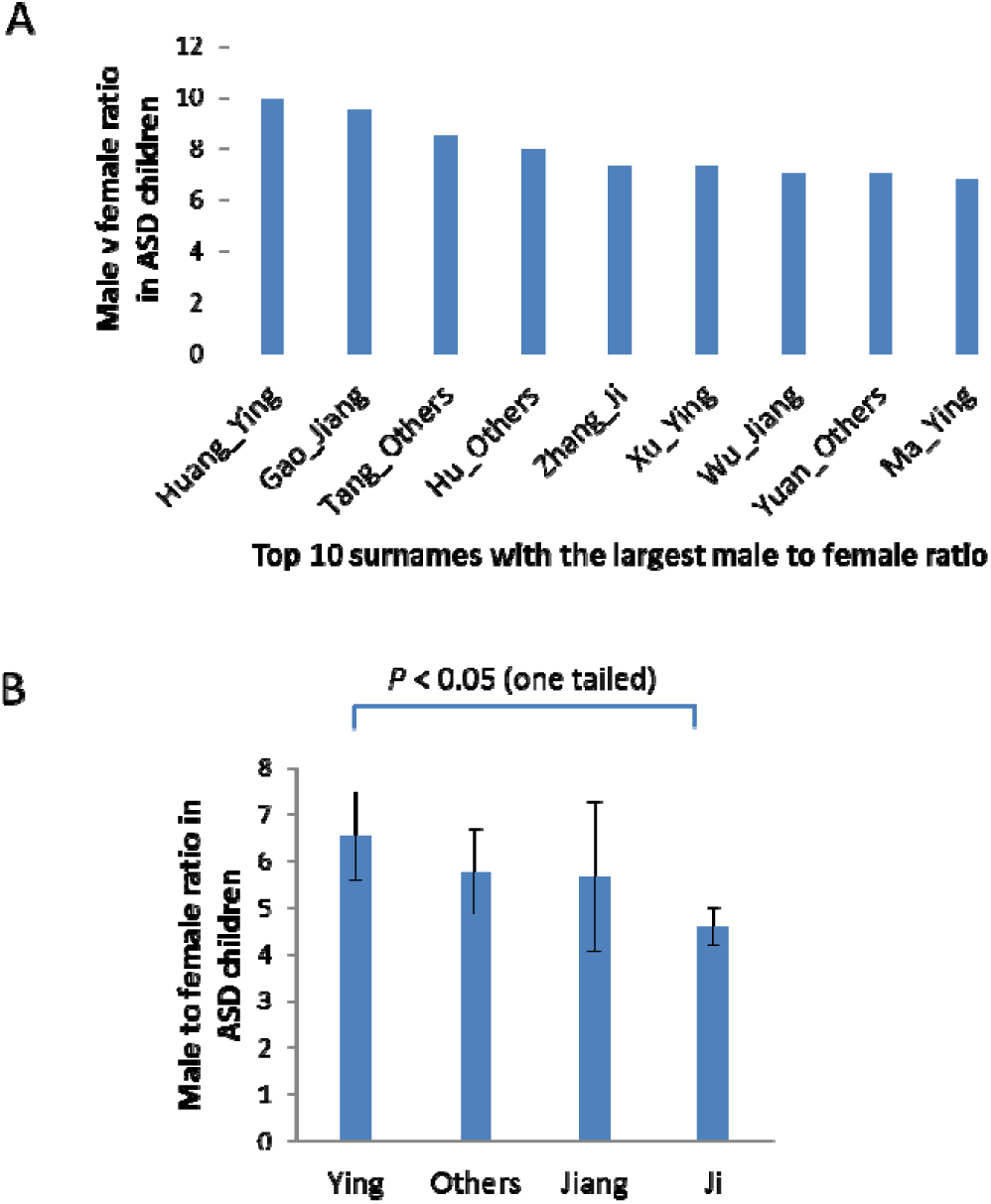
Male to female ratio in ASD children. Only top 40 surnames of China were counted among the 2362 ASD children studied. The minimal number of female cases was 2 in order to for a surname to be considered. **A.** Male to female ratio of ASD children with surnames among the top 10 most sex-biased surnames. **B.** Average male to female ratio of ASD children for each of the 4 groups of surnames. Also shown are the Standard Errors of the Mean and P value from Student’s t test, one tailed.

## Discussion

The association of the Jiang-group and Ying-group of surnames with O3a1c and O3a2c1a respectively suggests *Yan* and *Huang* as the candidate ancestors of these haplotypes, respectively. That Ji-group of surnames has less representation of O3a2c1a than Ying-group indicates more admixture for Ji related populations. Since the *Huang*-related haplotype O3a2c1a is the youngest among the three, and yet has claimed similar if not more descendants than the *Yan*-related O3a1c, the oldest of the three, the pace of expansion for the *Huang* lineage appears to be the fastest among the three Y haplotypes, consistent with *Huang* being the ultimate victor among the legendary leaders.

Previous studies on Europeans have produced either weak or no associations between Y haplotypes and ASD (*17, 18*). However, those studies did not consider surnames and finer classifications of Y haplotypes. The top ranked high SES descendant haplotype (178, 131, and 57) under each of the 3 super-grandfathers consistently showed a trend of lower fraction in ASD subjects with Ying related surnames, while the bottom ranked haplotypes (219, 161, 140, 40) showed the opposite. Also, ASD children of the Ying group showed expected higher male to female ratio. Therefore, population stratification is unlikely to account for these results.

Our results suggest a role for Y haplotypes in the male bias in ASD. The results also indicate that certain haplotypes may confer fitness advantages. A super-grandfather may leave many descendants for at least two reasons, fitter Y and more partners. Descendants with Y haplotypes more similar to the super-grandfather would continue to enjoy fitter traits and leave more descendants. Future studies along this line of investigation may help understand sex dimorphism in other diseases (*19*).

## Acknowledgements

Supported by the National Natural Science Foundation of China (81171880, 81330027, 81525007 and 31400919) and the National Basic Research Program of China (2011CB51001, 2012CB517900).

## Author contributions

SH and PH conceived the project. PH, NC, and ZZ performed DNA analysis. ZH and KX contributed the ASD and some normal DNA samples. SH and PH wrote the manuscript and all authors provided comments on the manuscript.

## Supplementary Information

### Supplementary Information text

1. Chinese surnames and the Eight Great Xings of High Antiquity
2. Supplementary Figure S1

**Supplementary Table S1.** Y chromosome haplotype distribution among Chinese surnames based on data from "One surname a week" website.

**Supplementary Table S2.** Distribution of the three Neolithic super-grandfather Y haplotypes among 772 male samples collected in this study.

**Supplementary Table S3.** Haplotype profiles of ASD and normal subjects.

**Supplementary Table S4.** Surname profiles of ASD children in the Autism Clinical and Genetic Resources in China.

**Supplementary Table S5.** List of SNPs for haplotype genotyping.

